# Applying high-resolution spatial transcriptomics to characterize the amyloid plaque cell niche in Alzheimer’s Disease

**DOI:** 10.1101/2023.06.29.546675

**Authors:** Anna Mallach, Magdalena Zielonka, Veerle van Lieshout, Yanru An, Jia Hui Khoo, Marisa Vanheusden, Wei-Ting Chen, Daan Moechars, I. Lorena Arancibia-Carcamo, Mark Fiers, Bart De Strooper

**Affiliations:** UK Dementia Research Institute at UCL, University College London, WC1E 6BT, London, UK; The Francis Crick Institute, London, NW1 1AT, UK; Department of Neurosciences and Leuven Brain Institute, KU Leuven, Leuven, Belgium; Laboratory for the Research of Neurodegenerative Diseases, VIB Center for Brain & Disease Research, VIB, Leuven, Belgium; BGI Research, BGI, Shenzhen, China; Discovery Biology, Muna Therapeutics, Leuven, Belgium; Department of Human Genetics, KU Leuven, Leuven, Belgium

## Abstract

The amyloid plaque cell niche is a pivotal hallmark of Alzheimer’s disease (AD). Where early spatial transcriptomics (ST) technologies have provided valuable information on transcriptomic alterations in the small tissue domains overlaying with amyloid plaques, they lacked cellular resolution. Here we compare two novel high-resolution ST platforms, CosMx and Stereo-seq, in their ability to characterize the cellular response in the amyloid plaque niche in an AD mouse model. Combining the results from both techniques empowered us to survey the highly variable microglial-astrocytic response across the amyloid plaque micro-environment and provided a first insight into how these responses could relate to neuronal transcriptomic alterations. This pilot study demonstrates the great potential of high-resolution ST, while simultaneously highlighting limitations that, when addressed, will unleash the full power of these techniques to map the progression of molecular and cellular changes in the brains of AD patients.

## Introduction

Alzheimer’s Disease (AD) is the most common form of dementia, characterized by progressive neurodegeneration that spreads throughout the brain. Widespread deposition of amyloid-β (Aβ) into plaques is a key neuropathological hallmark of the disease. Recent studies in the field have elucidated the importance of studying the so-called cellular phase of AD^1^, including how various cells within the brain respond to pathology. Changes in microglia and astrocytes, the support cells of the brain, have been strongly implicated in disease progression, with disease-associated microglia (DAM)^2–4^ and disease-associated astrocytes (DAA)^5–7^ identified as cell states of microglia and astrocytes that are strongly associated with AD.

We previously used spatial transcriptomics (ST), in conjunction with *in situ* sequencing, to identify a network of co-expressed genes, termed Plaque Induced Genes (PIGs), which co-localize with amyloid plaques and whose connectivity and expression levels increase with increasing Aβ load in *App^NL-G-F^* mice^8^. The microglial and astrocytic nature of the PIGs provides evidence for a spatially determined multicellular response to amyloid stress, involving the complement system, oxidative stress, lysosomes, and inflammation. The identification of this co-expression module suggests that microglia-astrocyte interaction strengthens with an increasing plaque load^8^. The low sensitivity of ST methodologies at that time, limited to 100 µm^2^ (Ref.^9^), hindered our ability to dissect specific cell type responses.

Since then, the rapid advancement of the field of ST has ushered in a number of platforms that boast cellular and subcellular resolution. Multiplexed Error-Robust FISH (MERFISH) is a single-molecule imaging method that uses combinatorial fluorescence *in situ* hybridization (FISH) to spatially profile 140-4,000 genes across 1 cm^2^ tissue sections^10,11^. The Xenium platform allows for the profiling of 300-400 genes across 2.8 cm^2^ (Ref^12^). Also belonging to the FISH class of methods, CosMx provides similar information across a total possible area of 3 cm^2^ (Ref^13^). Other methods, that are not commercially and therefore less widely available, are coppaFISH^14^ and STARmap PLUS^15^. All these technologies provide cellular resolution of a predetermined number of genes. On the other hand, sequencing-based techniques allow for *unbiased* profiling of the transcriptome. Slide-seq offers 10 µm spatial resolution on a 9.4 mm^2^ surface covered with DNA-barcoded beads with known positions^16,17^. Stereo-seq^18,19^ provides an enhanced 500 nm resolution in large-scale tissue slices (up to 1 cm^2^) by taking advantage of a DNA Nanoball (DNB) technology. In addition, the acquisition of a nuclear stain image of the section processed in Stereo-seq theoretically allows for the segmentation of transcripts into cells.

With the focus of comparing technologies that are commercially available and therefore feasible for institutes and core facilities to set up with ease, we decided to test CosMx, due to its large panel size of 950 genes^13^, and Stereo-seq, because of its unbiased detection of gene transcripts at unprecedented high-resolution^18,19^. We assessed these complementary methodologies in parallel to explore how their improved cellular resolutions can expand our knowledge about the PIG module^8^. Furthermore, we explored microglia and astrocyte interactions in AD and described the heterogeneity of the local cellular environment in the plaque niche, which is influenced by microglia-and astrocyte-driven glial responses. Ultimately, the two approaches provide complementary results, revealing the strength of combining both techniques to study these complex cellular interactions and downstream cellular effects on neurons. We also discuss some shortcomings which need to be considered when using these technologies to study molecular neuropathology of AD.

## Results

For both Stereo-seq and CosMx experiments, we used coronal sections from *App^NL-G-F^* and C57BL/6 mice^20^ (Fig 1A and D). This matches the brain regions and mouse models used in our previous ST study^8^.

**Figure 1:**
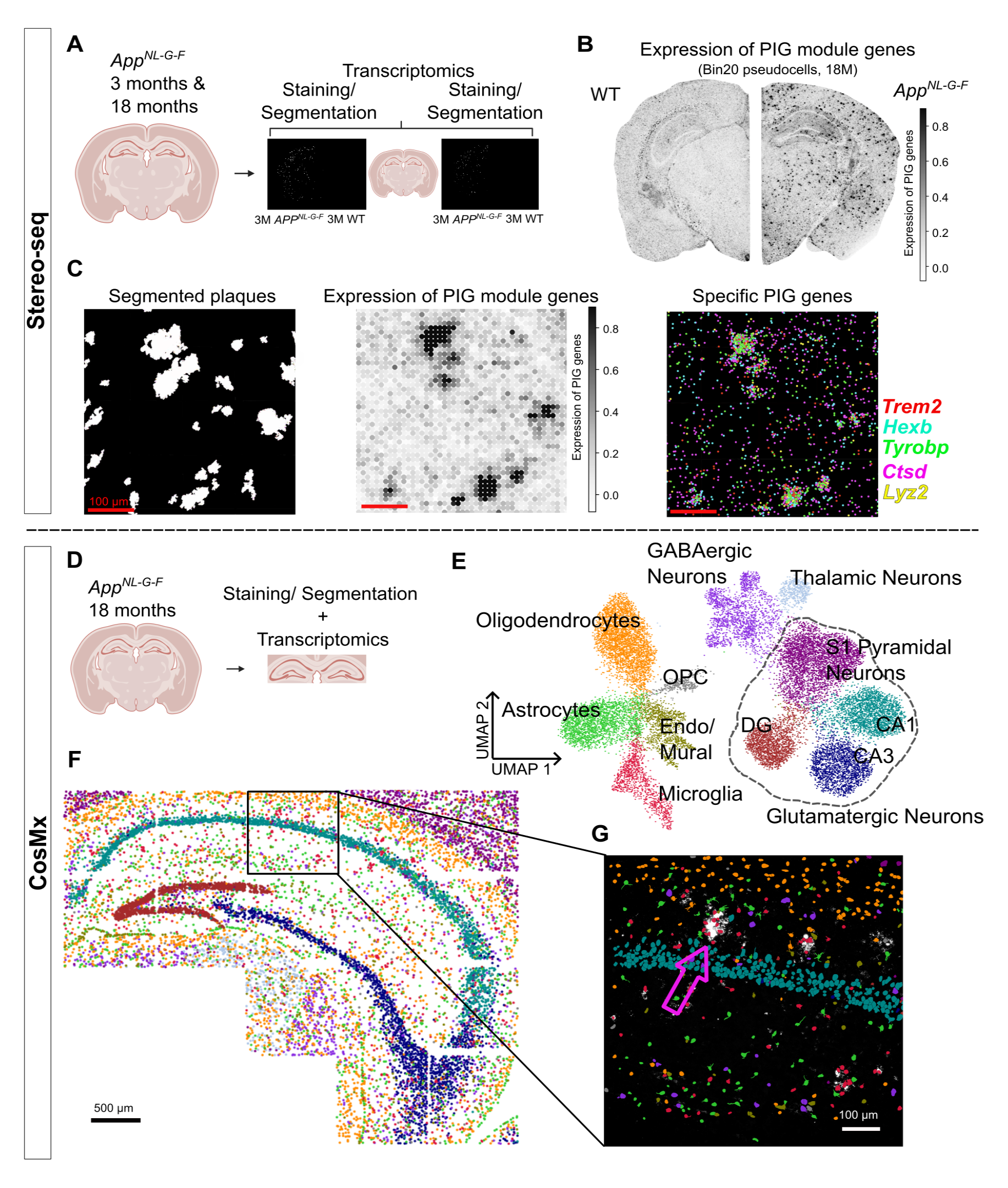
Resolving the plaque niche using spatial transcriptomics. A) Schematic of the experimental setup for Stereo-seq experiment. Two full coronal sections of 18-month (18M) *APP^NL-G-F^* mice and one full coronal section from an age-matched control were analyzed. For the 3-month (3M) timepoint, two half coronal sections were analyzed for the *APP^NL-G-F^* mice and two half coronal sections for the age-matched controls. Amyloid pathology was stained with the 6E10 antibody on adjacent slides and aligned with the transcriptome using the Fiji “Landmark correspondences” macro. B) Pseudo-cells from an 18M AD and an age-matched control hemisphere, scored and colored based on relative expression of the PIG module (white: low expression of PIG, black: high expression). Pseudo-cells in the AD hemisphere show marked elevation of PIG expression. Scores were obtained using the *score_genes()* function in SCANPY. C) Left: Representative image of segmented plaques in white from an 18M *App^NL-G-F^* mouse. Middle: pseudo-cells in the same coordinate space as above, scored on expression of PIG module (black: high expression of PIG). Right: spatial location of select PIG module transcripts (*Trem2, Hexb, Tyrobp, Ctsd, Lyz2*), also in the same coordinate space. D) Experimental setup for the CosMx experiments. The hippocampi and adjacent cortical regions from one coronal section from an 18M *App^NL-G-F^* mouse and from one age-matched control were assessed. Plaques were resolved in the same section using the monoclonal Aβ antibody MOAB-2. E) Uniform manifold approximation and projection (UMAP) showing the 20,024 segmented CosMx cells. Cells were unbiasedly clustered using the Leiden algorithm and annotated using a reference mouse brain dataset^23^. The glutamatergic neuron cluster was further resolved into sub-clusters, which were annotated based on spatial location. F) Spatial localization of the CosMx cells from the AD hemisphere, colored by Leiden cluster (as shown in E). Spatially resolving the clusters facilitated the annotation of the glutamatergic neuron subtypes. Scale bar represents 500 µm. G) Zoom-in image of segmented plaques (white), overlaid with segmented cells (colored by Leiden cluster, as shown in E and F) in the same coordinate space. One representative plaque was highlighted with the arrow. AD: *App^NL-G-F^* mice, WT: age-matched control, scale bar represents 100 µm, apart from F where the scale bar represents 500 µm.

For Stereo-seq, we obtained three adjacent coronal brain cryosections (each 10 μm apart) from one 3 month (3M) old *App^NL-G-F^* mouse and one age-matched control, as well as from two 18 month (18M) old *App^NL-G-F^* mice and one age-matched control (Fig 1A). The large capture area of Stereo-seq enabled us to profile the full coronal sections. The two outer sections were immunofluorescently stained for amyloid pathology using the 6E10 antibody, whereas the middle one was processed for Stereo-seq sequencing. Sequenced transcripts were binned into 20 x 20 DNB^18^ to obtain pseudo-cells with the size of 10 µm in width/height (Supplementary Fig 1E). We obtained between 445,039 and 512,367 pseudo-cells per chip, adding up to 2,425,202 pseudo-cell transcriptomic profiles across five chips. To assign each pseudo-cell to a region of the brain, we aligned each coronal section to a reference image from the Allen Mouse Brain Atlas^21^, identifying the white matter and 13 anatomical brain regions (Supplementary Fig 1A). Finally, we aligned the two adjacent sections from each sample to annotate each pseudo-cell with its respective distance to the nearest edge of plaque pathology.

In the CosMx platform, cell segmentation affords confident annotation of cell types (Fig 1E). We analyzed data from 0.15 cm^2^, equating to 27 fields of view (FOV), covering the hippocampus and small adjacent isocortical regions of coronal sections from an 18M *App^NL-G-F^* mouse and aged-matched control (Fig 1D). CosMx allows for protein staining directly on the same tissue section as the one used for transcriptomics, enabling direct annotation of cells with respect to their distance to plaque pathology. In addition to staining for amyloid-β using monoclonal Aβ antibody MOAB-2, the section was also stained for 18s RNA, histone, and GFAP to aid in the segmentation of transcripts into cells, following a previously described segmentation pipeline^13^.

### Cell identification in CosMx and use of pseudo-cells in Stereo seq

We were able to clearly identify the major expected cell types in the CosMx data set (Fig 1E and Supplementary Fig 1F). We note that the glutamatergic neuron cluster could be further separated into four subclusters, identified as S1 Pyramidal, CA1, CA3, and dentate gyrus (DG) neurons based on their spatial distribution within the tissue (Fig 1F).

In contrast, we were not able to recapitulate distinct cell type clusters and to confidently annotate cells with cell types using cell-segmented data based on nuclear staining^19^ in the much larger Stereo-seq data set (Supplementary Fig 1C and 1D). We hypothesize that the effect of RNA diffusion^18^, along with potentially suboptimal cell segmentation, prevented the generation of “clean” cells that could be confidently annotated. We therefore proceeded with downstream analyses using 20 x 20 DNB binned pseudo-cells as indicated (Supplementary Fig 1E).

To confirm that this binning approach has biological relevance^18^ we scored the Stereo-seq pseudo-cells^18^ on their expression of PIG module genes (Fig 1B). The pseudo-cells with high PIG expression superimpose over the plaque staining (Fig 1C). A correlation analysis between the mean PIG expression levels of pseudo-cells in function of their distance to plaques confirmed the higher expression of PIG module genes in close proximity to plaque pathology (R = -0.867, p = 9.80e-243) (Supplementary Fig 1B). We also assessed the spatial localization of specific PIG transcripts with respect to plaques and confirmed their overlap (Fig 1C). Thus, we can generate biological meaningful data using the pseudo-cells generated in the Stereo seq experiments^18^.

### The amyloid plaque cellular niche

We have previously shown the complex global cellular responses that reflect astro-and microgliosis and neuroinflammation in the amyloid plaque niche^8^. We did not have the resolution however to further dissect the contribution of individual cells and the cellular variation in different amyloid plaque niches.

We first explored the Stereo-seq dataset, relying on the cell2location algorithm^22^ to deconvolve the transcriptomes of plaque niches, defined here as the area within 40 µm of the plaque edge. We considered all transcripts within this area, inclusive of those overlaying the plaque, and deconvoluted them into cell types using a mouse brain reference dataset^23^ (Fig 2A and Supplementary Fig 2A). As the size of plaque niches varies, we compared cell type densities (the numbers of each cell type divided by the total number of cells present in a specific plaque niche) across plaque niches. A UMAP of the cell type compositions of plaque niche neighborhoods reveals a notable heterogeneity with respect to relative microglial densities ranging from 4% to 50% (Fig 2A), corresponding to a range of 0.3 to 20 estimated microglial cells per plaque niche (Supplementary Fig 2B). This heterogeneity in microglial densities can be observed in all analyzed brain regions (Fig 2A) and is particularly pronounced in plaques from 18M old mice (n=3,603 plaque niches; mean ± SEM: 19 % ± 0.1%; mean cell count ± SEM: 4.7 ± 0.04). This experiment also demonstrates that the increase in microglial cell density is related to the progression of amyloid pathology with age. The CosMx dataset is limited by the sample size (n=300 plaque niches), but clearly confirms the strong accumulation of microglia within the first 5-10 µm from the plaque edge (Fig 2C)^8,15,24^.

**Figure 2:**
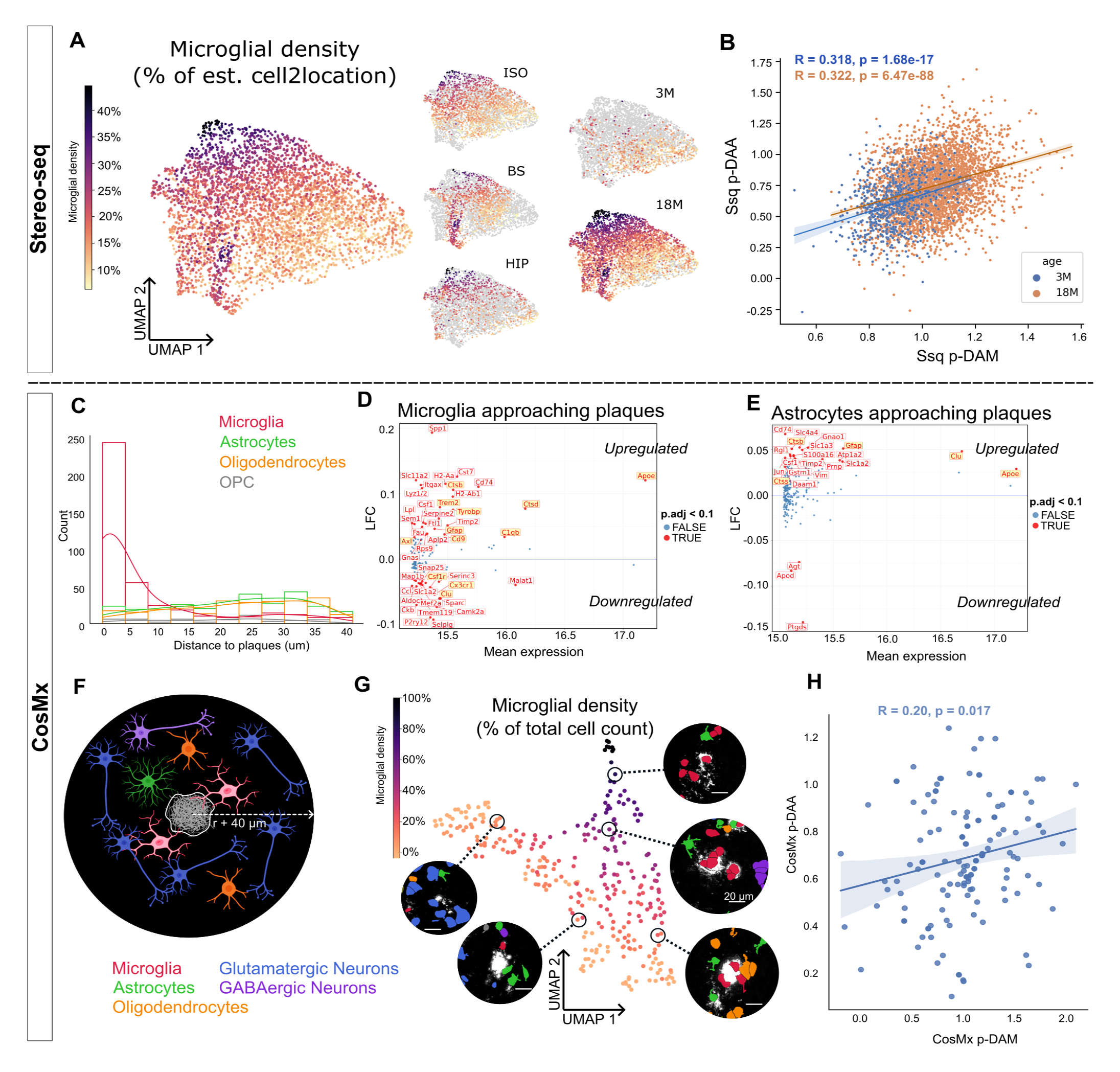
Cellular composition of the amyloid plaque niche. A) Left: The UMAP of the Stereo-seq plaque niches generated based on estimated cell type densities calculated using cell2location. Each point in the UMAP represents a plaque niche, colored by its estimated microglial density (calculated by dividing the number of estimated microglia by the total number of estimated cells in the plaque niche). Cell type densities were estimated using cell2location. Right: The same UMAP, with only plaque niches belonging to a certain region or to a certain age colored according to estimated microglial density. The remainder of the plaque niches are shown in gray. ISO: isocortex, BS: brainstem, HIP: hippocampus; 3M: 3 months, 18M: 18 months B) Pearson’s correlation between Ssq p-DAM (x-axis) and Ssq p-DAA (y-axis) levels in 3M (blue) and 18M (orange) plaque niches. Ssq p-DAM levels were defined as the expression level of DAM marker genes^3^ in the plaque niche minus the expression level of HM marker genes^3^ in the plaque niche. Ssq p-DAA levels were defined as the expression of DAA marker genes^5^ in the plaque niche minus the expression level of homeostatic astrocyte marker genes^5^ in the plaque niche. C) Histogram and density plots of cell type distributions (y-axis) at increasing distances from plaques (x-axis), as seen in the CosMx dataset. D) MA plot showing gene differential expression in microglial cells with respect to square root of distance to plaques, with the x-axis showing mean expression and the y-axis log fold change (LFC). Genes upregulated in cells close to plaques are shown above the blue line (y = 0). Quasi-likelihood F-test (QLFTest), P-values adjusted with Benjamini-Hochberg correction (P.adj < 0.1). Significantly changed genes are labeled in red. Genes belonging to the PIG module are additionally highlighted in yellow. E) MA plot showing gene differential expression in astrocytes with respect to square root of distance to plaques. Genes upregulated in cells close to plaques are shown above the blue line (y = 0). Quasi-likelihood F-test (QLFTest), P-values adjusted with Benjamini-Hochberg correction (P.adj < 0.1). Significantly changed genes are labeled in red. Genes belonging to the PIG module are additionally highlighted in yellow. F) A representative diagram highlighting the average cellular composition of a plaque niche. The plaque niche is defined as a circle with center equal to the plaque niche’s centroid and radius (r) equal to the plaque niche’s radius + 40 µm. The average amyloid plaque cellular niche consists of two microglia (red), two oligodendrocytes (orange) and one astrocyte (green) besides one GABAergic neuron (purple) and five glutamatergic neurons (blue). Sizes of the different cell types not to scale. Please note that the cellular composition of the amyloid plaque niche varies strongly as discussed below. G) The UMAP of the CosMx plaque niches was generated based on the cell type densities in the neighborhood of each plaque. Each point represents a plaque niche, colored by its estimated microglial density. Representative plaque niches and the cellular composition of their neighborhoods are shown, with segmented plaques in white and segmented cells colored according to cell type (as outlined in F). Scale bar represents 20 µm. H) Pearson’s correlation between CosMx p-DAM level (x-axis) and CosMx p-DAA level (y-axis) of each plaque niche. CosMx p-DAM level is defined by the expression of DAM markers of the microglia belonging to a plaque niche, whilst CosMx p-DAA level is defined by the expression of DAA markers of astrocytes belonging to a plaque niche.

The question arises whether the transcriptome of the cells in the amyloid plaque niche is indeed different from the cells outside of this niche. We therefore investigated the changes in the transcriptomes of specific cell types as they approach plaque pathology (Figure 2D, 2E and Supplementary Fig 2C). We confirm notable changes in the microglial transcriptome (28 genes upregulated and 29 genes downregulated). Marker genes for homeostatic microglia (HM), such as *Cx3cr1* and *Tmem119*, are downregulated in microglia as they approach plaques (Fig 2D). On the other hand, disease-associated microglia (DAM) markers^2,3^, including *Apoe, Lpl* and *Trem2*, are upregulated in proximity to plaques.

Interestingly, Disease Associated Astrocyte (DAA) genes, such as *Gfap, Ctsb* and *Vim,* are also significantly upregulated in astrocytes as they approach plaques (Fig 2E) indicating that the amyloid plaques also drive astrocytic pathological cell states. A direct comparison of the microglial and astrocytic responses to pathology revealed that genes linked to the PIG module (Fig 2D and 2E, highlighted in yellow) are predominantly changed in microglia (Supplementary Fig 2D), which remained a matter of speculation using lower resolution ST methodologies^8^.

We next narrowed in on the plaque niches in this CosMx dataset, likewise defined as all cells located within 40 µm from a plaque edge, to further analyze the cellular composition of the amyloid plaque niche. We find that the average cellular composition of a CosMx plaque niche (represented in Fig 2F) contains 2 microglia, 1 astrocyte, 2 oligodendrocytes, 1 GABAergic and 5 glutamatergic neurons. The cellular composition of amyloid plaques is highly heterogeneous (Fig 2G, with images representing real plaque niches observed in the sample, and Supplementary Fig 2E).

We wondered to what extent AD-associated cell state signatures of microglia and astrocytes were related (which would suggest cellular crosstalk). We assigned a DAM score^3^ to each plaque niche, which we named p-DAM. In the CosMx dataset, this value is in essence based on the expression level of DAM genes in the microglia (referred to as CosMx p-DAM). In the Stereo-seq we calculated the DAM score of each plaque niche (referred to as Ssq p-DAM) by calculating the module score of DAM genes relative to the expression of HM genes. This corrects for the fact that the HM module score will also increase when the microglial numbers go up in the plaque niche (Supplementary Fig 2G). Following similar approaches, we also assigned each plaque an astrocyte relevant DAA level^5^ (p-DAA) in both datasets. The Ssq p-DAM and Ssq p-DAA levels of plaque niches are strongly correlated (Fig 2B), supporting the notion of microglial-astrocytic crosstalk in close proximity to plaques^8^. We noted a similar, albeit weaker, correlation between CosMx p-DAM and CosMx p-DAA (Fig 2H). As there are fewer plaque niches analyzed in the CosMx data set, the correlation is much less confident than that seen in Stereo-seq (compare Fig 2B and 2H).

In conclusion, these experiments show that the amyloid plaques in this model recruit specific cellular cell states, which apparently mutually influence each other.

### Brain region-specific aspects of the glial responses in the amyloid plaque cell niche

We wondered to what extent the variations in cells and cell states were driven by brain region. Unbiased clustering of the Stereo-seq transcriptomes around the amyloid plaques (Fig 3A) demonstrates that the transcriptomic heterogeneity in plaque niches is mainly driven by the type of neurons characteristic for the brain area in which the plaque is found (Supplementary Fig 3E). That said, DAM genes are consistently and similarly upregulated in pseudo-cells in the proximity of plaques, independent of whether they are located in the isocortex or hippocampus (Fig 3B, r_s_ = 0.817, p < 2.2e-16). The DAA genes finally show greater regional variability (Fig 3C, r_s_ = 0.583, p < 2.2e-16), indicating that the transcriptomic responses of astrocytes to plaque pathology, i.e. p-DAA, are more variable across regions.

**Figure 3:**
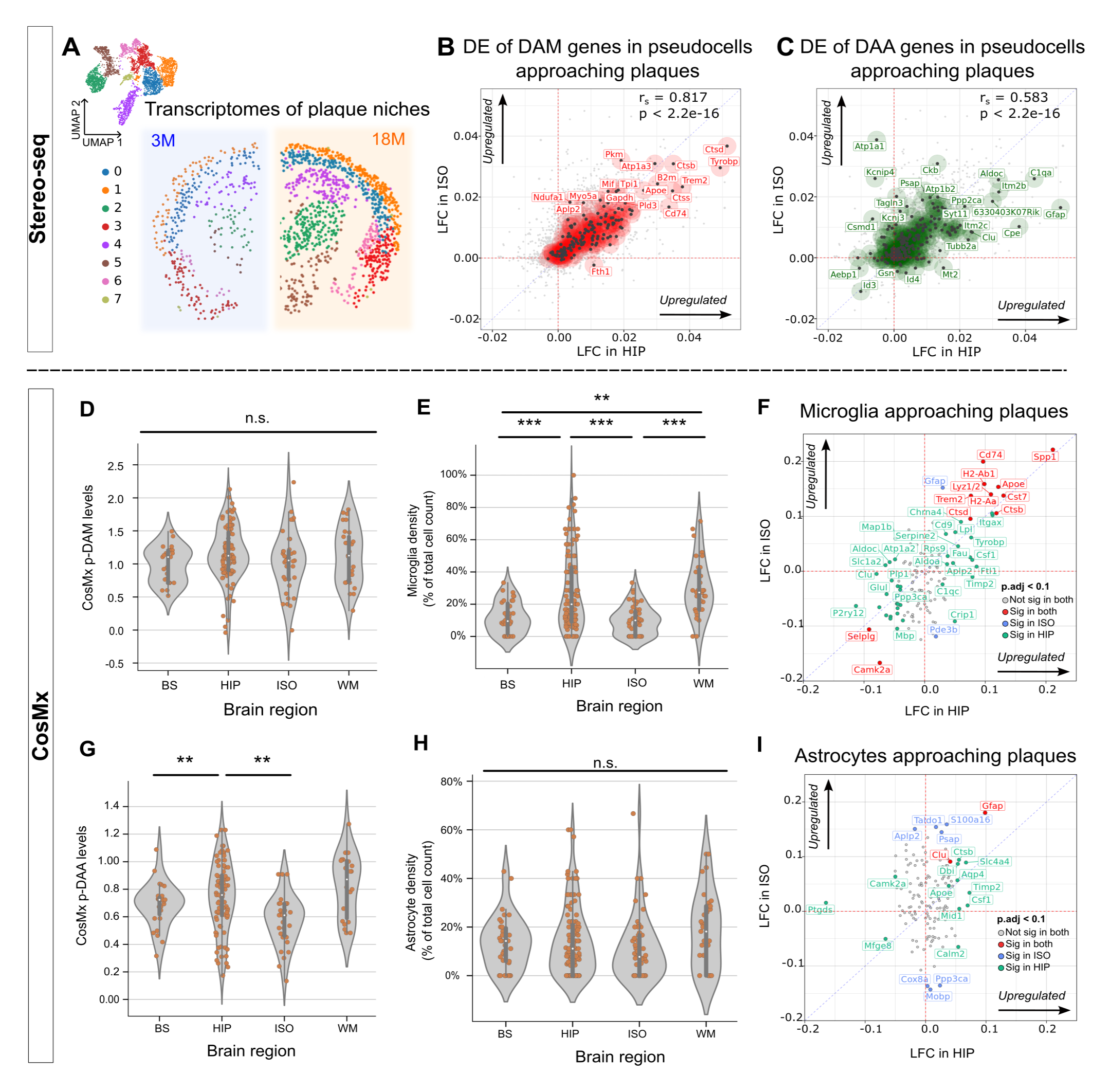
Brain region-specific variations of the plaque niche. A) UMAP representing the integrated and unbiasedly clustered transcriptomes of Stereo-seq plaque niches from all 18M and 3M *App^NL-G-F^* mice and the spatial distribution of the clusters in a 3M and 18M hemisphere. Plaque niches are unbiasedly clustered using the Leiden algorithm. B) Quadrant plot showing the LFCs of genes in pseudo-cells with respect to distance to pathology in the hippocampus (x-axis) and isocortex (y-axis). Red: DAM marker genes. LFCs were calculated using a Quasi-likelihood F-test (QLFTest). r_s_ values were calculated using Spearman’s correlation. C) Quadrant plot showing the LFCs of genes in pseudo-cells with respect to distance to pathology in the hippocampus (x-axis) and isocortex (y-axis). Green: DAA marker genes. LFCs were calculated using a Quasi-likelihood F-test (QLFTest). r_s_ values were calculated using Spearman’s correlation. D) Violin plots of CosMx p-DAM levels across different regions in the CosMx dataset. No significant differences between regions were detected. One-way ANOVA followed by Tukey HSD multiple comparisons correction. E) Violin plots comparing microglial densities (percentage of microglia in relation to total number of cells in the plaque niche) of plaque niches across different brain regions in CosMx. One-way ANOVA followed by Tukey HSD multiple comparisons correction. F) Quadrant plot showing the similar transcriptomic responses of microglia to plaque pathology in the isocortex (y-axis) and in the hippocampus (x-axis) in the CosMx dataset. Red: genes significant in both comparisons, gray: genes not significant in either comparison, blue: genes significant in isocortical microglia only, green: genes significant in hippocampal microglia only. Quasi-likelihood F-test (QLFTest), P-values adjusted with Benjamini-Hochberg correction (P.adj < 0.1). G) Violin plots comparing CosMx p-DAA levels across different brain regions. Region-specific differences were observed. One-way ANOVA followed by Tukey HSD multiple comparisons correction. H) Violin plots comparing astrocytic densities (percentage of astrocytes in relation to total number of cells in the plaque niche) of plaque niches across different brain regions in CosMx. One-way ANOVA followed by Tukey HSD multiple comparisons correction. I) Quadrant plot comparing the transcriptomic responses of astrocytes to plaque pathology in the isocortex (y-axis) and in the hippocampus (x-axis). A stronger response of astrocytes is seen in the isocortex. Red: genes significant in both comparisons, gray: genes not significant in either comparison, blue: genes significant in cortical astrocytes only, green: genes significant in hippocampal astrocytes only. Quasi-likelihood F-test (QLFTest), P-values adjusted with Benjamini-Hochberg correction (P.adj < 0.1). WM: white matter, ISO: isocortex, HIP: hippocampus, BS: brainstem; *P < 0.1; **P < 0.01; ***P < 0.001; ns, not significant.

We further investigated the regional dependency of p-DAM and p-DAA in CosMx, where we could take advantage of the cell segmentation. Despite regional differences in microglia densities (Fig 3E) and raw microglia numbers (Supplementary Fig 3A), we did not detect differences in CosMx p-DAM levels across different brain regions (Fig 3D and 3F). On the other hand, we observed regional variations in CosMx p-DAA levels (Fig 3G), independent of astrocyte densities (Fig 3H) and raw cell counts (Supplementary Fig 3B) which were consistent across brain regions. Thus, the CosMx and the Stereo-seq data indicate that astrocytic DAA levels in the amyloid plaque niches vary over the different brain areas. Furthermore, we observed that DAA-related genes, such as *Ctsb*, are only significantly differentially expressed in hippocampal astrocytes, but not in isocortical astrocytes, as they near plaques (Fig 3I). Based on these findings (Fig 3C and 3I), we propose region-specific differences in astrocytic response to plaques. The DAA phenotype might mainly capture the response of hippocampal astrocytes, where this phenotype was originally identified^5^.

### Neuronal responses to the glial changes in the amyloid plaque cell niche

To explore the effects of the microglia and astroglia on the other cells in the amyloid plaque cell niche, we investigated differentially expressed genes in response to Ssq p-DAM (Fig 4A) and Ssq p-DAA (Fig 4B). Interestingly, these transcriptomic changes increase strongly from 3M to 18M. For the DAM response, we see 346 genes significantly differentially expressed at 3M (Supplementary Fig 4A) compared to 2,430 genes at 18M (Fig 4A) while for the DAA response 145 genes are differentially expressed at 3M (Supplementary Fig 4B) in comparison to 668 genes at 18M (Fig 4B). Out of the genes that change in function of Ssq p-DAM and Ssq p-DAA at 18M, 75% of them are up-regulated or down-regulated in *both*, suggesting a shared response of cells in the cell niche to the micro-and astroglia alterations (Supplementary Fig 4C). We performed gene set enrichment analyses (GSEA) with respect to the Ssq p-DAM and Ssq p-DAA axes (Supplementary Table 1), visualized in the quadrant plot in Fig 4C.

**Figure 4:**
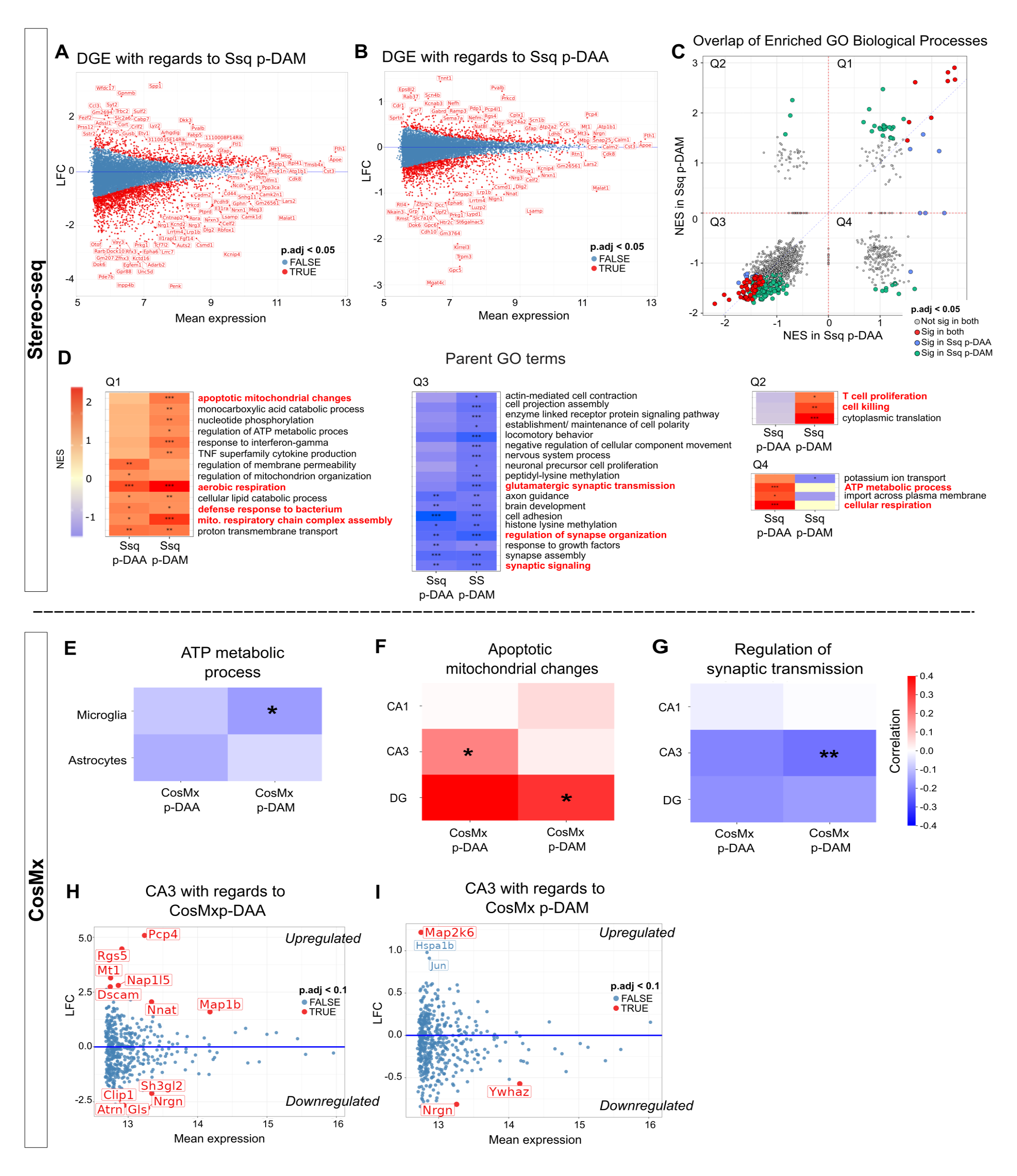
Effect of p-DAM and p-DAA on predicted cellular functions. A) MA plot showing gene differential expression in 18M Stereo-seq plaque niches (pseudo-bulked) with respect to the niche Ssq p-DAM level. Genes upregulated in plaque niches with higher p-DAM levels are shown above the blue line (y = 0). Significantly changed genes are labeled in red. Quasi-likelihood F-test (QLFTest), P-values adjusted with Benjamini-Hochberg correction (P.adj < 0.05). B) MA plot showing gene differential expression in 18M Stereo-seq plaque niches (pseudo-bulked) with respect to the niche Ssq p-DAA level. Quasi-likelihood F-test (QLFTest), P-values adjusted with Benjamini-Hochberg correction (P.adj < 0.05). C) Quadrant plot comparing normalized enrichment scores (NES) for GO biological processes (BP) terms in 18M plaque niches in response to Ssq p-DAM levels (y-axis) and to Ssq p-DAA levels (x-axis) (Supplementary Table 1). Normalized Enrichment Scores (NES) were obtained by gene set enrichment analyses (GSEAs). The vast majority of significantly enriched terms (321 out of 336 in total) are present in either quadrant 1 (Q1) or Q3 (i.e. they are significantly enriched in response to both p-DAM and p-DAA levels). Red: terms significantly enriched in both, gray: terms not significant in both, blue: term significantly enriched in p-DAA, green: terms significantly enriched in p-DAM. D) Heatmaps summarizing the NES (color scale) and significance of enriched biological processes terms from each quadrant (as shown in A). Significant terms from each quadrant were aggregated into parent terms, and the average NES scores of the terms for both the p-DAM and p-DAA GSEAs were plotted. Terms of interest are highlighted in red. E) Pearson’s correlation between the expression levels of genes linked to the “ATP metabolic process” GO term, and p-DAM and p-DAA levels in the CosMx dataset for microglia and astrocytes (R value, color scale). Genes linked to this biological function were negatively correlated with increasing CosMx p-DAM in microglia. F) Pearson’s correlation between the expression levels of genes linked to the “Apoptotic mitochondrial changes” GO term, and p-DAM and p-DAA levels in the CosMx dataset for different subclasses of glutamatergic neurons (R value, color scale). G) Pearson’s correlation between the expression levels of genes linked to the “Regulation of synaptic transmission” GO term, and p-DAM and p-DAA levels in the CosMx dataset for different subclasses of glutamatergic neurons (R value, color scale). Genes linked to this biological function were negatively correlated with increasing p-DAM levels in CA3 neurons. H) MA plot showing gene differential expression in CA3 neurons in a plaque niche with respect to CosMx p-DAA levels. Significantly changed genes are labeled in red. Quasi-likelihood F-test (QLFTest), P-values adjusted with Benjamini-Hochberg correction (P.adj < 0.1). I) MA plot showing gene differential expression in CA3 neurons in a plaque niche with respect to CosMx p-DAM levels. Significantly changed genes are labeled in red. Quasi-likelihood F-test (QLFTest), P-values adjusted with Benjamini-Hochberg correction (P.adj < 0.1). *P < 0.1; **P < 0.01; ***P < 0.001

In line with the above, there is a large overlap between significantly enriched gene ontology (GO) biological processes are either up-or down-regulated in response to both Ssq p-DAM *and* Ssq p-DAA. This is reflected in their location in quadrant 1 (Q1) or Q3, respectively. Further analysis of the enriched terms, as well as their enrichment patterns (Fig 4D), indicate that terms linked to inflammation, such as “defense response to bacterium”, but also “TNF superfamily cytokine production” or “response to interferon-gamma” are linked to plaque niches with high Ssq p-DAM or Ssq p-DAA levels. Likewise, GO terms linked to energy production (such as “aerobic respiration” and “mitochondrial respiratory chain complex assembly”) are upregulated in higher Ssq p-DAM and Ssq p-DAA plaque niches. Whilst we lack the cellular resolution in Stereo-seq to identify the cell types driving these changes, microglia are known to increase their energy production to launch an inflammatory response under stress^25,26^. Meanwhile, we saw a downregulation of genes linked to neuronal function, such as “synaptic signaling/transmission” and “regulation of synapse organization" (Fig 4D) with respect to both Ssq p-DAM and Ssq p-DAA. This highlights the potential detrimental effect of specific plaques with high p-DAM and p-DAA levels on neuronal functioning.

Although the cellular responses to Ssq p-DAA and Ssq p-DAM appear to be largely consistent, a closer look at the biological terms located in Q2 and Q4 reveals that certain biological terms are only enriched in response to either Ssq p-DAM or Ssq p-DAA (Fig 4D). For example, *Traf3*, a component of the Tumor Necrosis Factor (TNF) receptor involved in neuronal death, inflammation, and oxidative stress in neurons^27,28^ (Supplementary Fig 4C), and the linked GO functions of the gene (Fig 4D, Q4) are specifically associated with increased Ssq p-DAM. On the other hand, genes such as *Atp2b1,* which is involved in the maintenance of intracellular calcium^29,30^, and their associated GO terms (Fig 4D, Q2) are associated with higher Ssq p-DAA. This indicates that whilst p-DAM and p-DAA appear to induce overlapping responses, certain pathways, such as those relating to cell killing and calcium signaling, may be specifically activated by one of the two axes.

Next, we sought to associate the proposed enriched GO terms from the Stereo-seq data to specific cell types using the CosMx data. Genes related to the GO terms “ATP metabolic process” are negatively correlated with increased CosMx p-DAM in microglia (Fig 4E). Also, apoptotic mitochondrial changes are significantly correlated with p-DAM and p-DAA levels in the DG and CA3 neurons, respectively (Fig 4F). The CA3 neurons show a significant negative correlation between their expression of “regulation of synaptic transmission” genes and their plaque niches’ CosMx p-DAM levels (Fig 4G). In line with this, direct analyses of CA3 neurons with respect to their p-DAM and their p-DAA levels reveals a significant downregulation of *Nrgn* (neurogranin), which is involved in long-term potentiation (LTP) signaling in neurons^31^, in both comparisons (Fig 4H/I, Supplementary Fig 4D-G).

## Discussion

In this study, we compared two novel high-resolution ST techniques to study the local changes around amyloid-β plaques in a mouse model of AD. The Stereo-seq and CosMx techniques were chosen for this study because of their similarly high subcellular resolution and commercial availability. We found that the data from both techniques was highly complementary (Table 1) and, when used in tandem, empowered us to answer important questions regarding the characterization of the amyloid plaque cell niches and how cell neighborhoods influence each other.

**Table 1:** Comparison of the Stereo-seq and CosMx experiments on key statistics. Here we compare the performance of Stereo-seq and CosMx on key metrics.

|  | Stereo-seq | CosMx |
| --- | --- | --- |
| Tissue area analyzed | Full coronal section, up to 0.9 cm<sup>2</sup> | 0.15 cm<sup>2</sup> |
| Number of genes | 12,535 | 950 |
| Plaque staining | On adjacent sections 10 μm away | On the same section |
| Clean cell segmentation? | No | Yes |

The PIG response has been characterized in several models^8,15,32^. Not only did we confirm that PIGs increase in expression in microglia and astrocytes near plaques (Fig 2D and 2E), we could also clarify that the majority of these previously-described gene changes happen in microglia rather than in astrocytes (Supplementary Fig 2D). Interestingly, two previously reported PIGs, *Csf1r* and *Cx3cr1*, are in fact downregulated in microglia as they approach plaques (Fig 2D). These genes are known markers for HM^3^, and thus their downregulation in proximity to amyloid pathology should be expected. This highlights the limitations of lower resolution ST methods, such as the one originally used to identify the PIG module^8^. As the number of microglia around plaques is markedly higher (Fig 2C), even homeostatic microglial markers can appear to be associated with plaques if the spatial resolution does not allow for cell type-specific analysis.

The PIGs describe a proposed microglia-astrocyte interaction. Our publication here and other recent publications using Visium^32^ or STARmap PLUS^15^ have begun to investigate the link between the PIGs and cell states defined by single cell/nucleus transcriptomics, revealing that microglia and astrocytes accumulating around plaques adopt a DAM and DAA phenotype respectively.

It appears that the microglial response in the plaque niche (defined here as p-DAM), is a universal response that is consistent across brain regions (Fig 3B, 3D and 3F), confirming previous work that DAM develops independent of the hippocampus or cortex region^3^. On the other hand, we observe that astrocytes in the hippocampus seem more prone to develop a DAA signature than in other regions (Fig 3C and 3I), as shown in the Stereo-seq (Supplementary Fig 3D) and CosMx (Fig 3G) data sets. Other studies have shown that astrocytes display region-specific differences, both in terms of their phenotype^33^ and their functions such as calcium signaling^34^. Furthermore, it is also known that hippocampal astrocytes develop more pronounced changes to aging, akin to developing a DAA phenotype, compared to isocortical astrocytes^35^. As the DAA signature was specifically characterized in hippocampal astrocytes^5^, it is possible that astrocytes in other regions of the brain might respond differently to plaques. Indeed, since the description of the DAA phenotype^5^, work from others has expanded the nomenclature of astrocyte substates^36,37^. Unfortunately, diving into this question further and describing the astrocytic response to plaques in different brain regions was outside of the scope of our current analysis, as we were limited to a panel of 950 genes in CosMx and lacked the cellular resolution to do so in Stereo-seq.

Going beyond the description of DAM and DAA around the plaques, we explored whether we could make predictions of the effect of these glial responses on the function of neurons, something which has not been done before in ST studies^15,32^. Focusing our analysis within the amyloid plaque cell niches (Fig 4) we observed that genes linked to synaptic transmission are downregulated in plaque environments correlating with elevated Ssq p-DAM and p-DAA levels (Fig 4D), an effect that was specifically clear in the hippocampal neurons, particularly CA3 neurons (Fig 4F/G). Whilst the *App^NL-G-F^* mouse model we used for this study does not show widespread neuronal loss^20^, previous research has suggested that synaptic function and long-term potentiation formation is impaired in this mouse model^20,38^.

We also found that cells display changes in their energy metabolism in plaque niches with elevated Ssq p-DAM and Ssq p-DAA (Fig 4D). Microglia change their energy production from oxidative phosphorylation to glycolysis to launch an inflammatory response^25,26^. In line with this, microglia show a downregulation of ATP metabolic process in response to CosMx p-DAM (Fig 4E). DAA display genes linked to increased energy production, especially lipid metabolism^5^, which we also link to increased Ssq p-DAA (and Ssq p-DAM) levels (Fig 4D). We show that Ssq p-DAM and Ssq p-DAA levels correlate, occurring at increased levels around the same plaques (Fig 2B), and that they induce similar transcriptomic and functional effects on other cells in their neighborhood (Supplementary Fig 4C, Fig 4C). However, to show that p-DAM does indeed drive p-DAA in a sequential manner, we would need to analyze a bigger dataset with more timepoints covering the gradual development of both responses. Previous literature suggests however also that microglial DAM-like responses can drive astrocytes to develop a toxic phenotype^6,39^

### Limitations/Next steps

Using Stereo-seq and CosMx as orthogonal methods in this study proved to be effective in overcoming the limitations of either technique. As we show, the lack of reliable cell segmentation is a serious limitation of Stereo-seq (Supplementary Fig 1C and 1D). A confounding factor is the diffusion of transcripts, which should be addressed in subsequent versions of this technique. The main shortcoming of the CosMx technology is the limited panel size. Furthermore, as with all high-resolution ST methodologies, accurate analysis on a cellular level is dependent on accurate segmentation of transcripts into cells. We note that highly expressed transcripts in the CosMx data are sometimes mis-segmented. For example, while we see in line with previous literature^5,40,41^ an upregulation of *Gfap* in astrocytes close to plaques (Fig 2E), we also see an upregulation of *Gfap* close to plaques in other cell types (Supplementary Fig 2C and 2D) that are not known to express these transcripts. A similar problem was noticed for *Apoe,* another transcript that is specifically enriched in microglia around plaques.

Overall while both novel spatial techniques appear useful and can be upscaled, further development of data analysis resources and improvements along the lines we suggest above are needed to maximize the output of these approaches. Ultimately, cells exist in 3D neighborhoods, and recent work has demonstrated very impressively that much can be learned from a 3D spatial analysis^42^. In the future, reduced costs of running these experiments would provide researchers with the ability to process a larger number of samples, which in turn will allow for the screening of more intermediate timepoints and to increase statistical power that enables the identification of less abundant events likely relevant to the disease process. Altogether, spatial analysis of single cells leads to a better understanding of cellular neighborhoods and how cell-cell interactions maintain homeostasis and how this might collapse in AD.

## Conclusion

In this study, we harnessed the power of two novel ST methods, combined with refined computational approaches, to describe in a proof-of-concept manner the cellular and transcriptomic variations in local plaque environments. We leveraged the strengths of both technologies, including the reliable cell segmentation and single-cell resolution provided by the CosMx platform as well as the unbiased transcriptomic profiling and large capture area of Stereo-seq, to expand on previously defined cell type signatures in plaque niches and to provide further evidence for microglial-astrocytic crosstalk in the niches. Furthermore, we were able to characterize the disruptive effects microglial and astrocytic activation levels have on neuronal functions in a model that does not show widespread neuronal loss.

## Author contributions

Conceptualization A.M., M.Z., M.V., W.-T.C., M.F., I.L.A.-C., B.D.S.

Methodology V.v.L., Y.A., J.H.K.

Software M.Z., D.M., M.F.

Validation A.M., M.Z., V.v.L.

Formal Analysis A.M, M.Z., V.v.L., D.M., M.F.

Investigation V.v.L, Y.A., J.H.K.

Resources A.M., M.V., W.-T.C., Y.A., J.H.K..

Data Curation M.Z., M.F.

Writing - Original Draft A.M., M.Z., V.v.L.

Writing - Review & Editing A.M., M.Z., I.L.A.-C., M.F., B.D.S.

Visualization A.M., M.Z.

Supervision A.M., M.F., L.A.-C., B.D.S.

Project administration A.M., M.F, I.L.A.-C., B.D.S.

Funding Acquisition B.D.S.

## Acknowledgements

We would like to thank BGI for providing us with access and training to the Stereo-seq platform and for the continued collaboration, as well as VIB Technology Watch member, Yu-Chun Wang, for facilitating collaborations with BGI research. We would like to thank Nanostring for making the CosMx experiment possible through the Technology Access Program and the Francis Crick Institute for funding this experiment. Thanks also goes to Tancredi Massimo Pentimalli, who inspired us to pursue neighborhood analyses by presenting his own CosMx analysis.

## Conflict of interests

B.D.S. has been a consultant for Eli Lilly, Biogen, Janssen Pharmaceutica, Eisai, AbbVie and other companies and is now consultant to Muna Therapeutics. B.D.S is a scientific founder of Augustine Therapeutics and a scientific founder and stockholder of Muna Therapeutics. M.F. is a consultant to Muna Therapeutics. Y.A. and J.H.K. are employees of BGI Research and M.V. and W.T.-C. are employed by Muna Therapeutics. W.T.-C. is also a stockholder of Muna Therapeutics.

*Supplementary Figure 1:*

A) Left: Alignment of brain regions as defined by the reference atlas from Allen Brain Institute to the Stereo-seq transcriptomic data. Right: pseudo-cells retained after QC and removal of white matter regions, colored by their respective region annotation.

B) Mean PIG expression levels of pseudo-cells (y-axis) at a given distance from plaques (x-axis), within 400 µm of plaque edge. Pearson’s correlation.

C) Left: UMAP of the cell-segmented Stereo-seq data, clustered using the Leiden algorithm. Right: Cell segmented cells annotated by relative expression of microglial marker genes. Cells with a high microglial score corresponded to cluster 7.

D) Left: UMAP of segmented cells (candidate microglia cluster) from cluster 7 (as shown in B), re-clustered using the Leiden algorithm. Right: top marker genes for each sub-cluster, obtained using the Wilcoxon rank sum test. Cells in this cluster contain neuronal, astrocytic, and oligodendrocytic marker genes in addition to microglial ones, instead of different microglia cell states as expected^3^, highlighting that the cell segmentation did not lead to well-segmented cells.

E) UMAPs of the pseudo-cell transcriptomes from all processed Stereo-seq chips, colored by brain region (left), age (middle), and phenotype (right). The differences in the transcriptomes are predominantly driven by region, with good integration between ages and genotypes.

F) UMAP of the cell-segmented CosMx data showing the expression level of cell type markers for each cell. Cell type markers were obtained from the Allen Brain Institute reference atlas (Yao 2021).

G) Spatial distribution of the segmented CosMx cells, annotated by respective brain regions.

*Supplementary Figure 2:*

A) UMAP of plaque niche cellular environments in Stereo-seq AD samples (3M and 18M integrated), colored by estimated cell type densities, as calculated by the total number of cells predicted in the plaque niche. The number of each of the different cell types in each plaque niche was estimated using cell2location.

B) UMAP from A, showing the predicted microglial numbers for each plaque niche, as calculated by cell2location, without adjusting for varying total number of cells in a plaque niche.

C) Based on the CosMx dataset, cell type specific MA plots showing gene differential expression in each cell type as they approach plaques. Genes upregulated in cells close to plaques are shown above the blue line (y = 0). Quasi-likelihood F-test (QLFTest), P-values adjusted with Benjamini-Hochberg correction (P.adj < 0.1). Significantly changed genes are labeled in red.

D) A comparison of log fold changes of PIGs in microglia (blue) vs. in astrocytes (orange) with respect to square root of distance to plaques. Positive LFCs indicate upregulation in cells close to plaques. The genes are colored by the cell types the previous publication^8^ predicted them to be expressed in, with the blue and orange genes proposed to be expressed in microglia and the green were proposed to be expressed in astrocytes.

E) UMAP of plaque niche cellular environments in CosMx AD hemisphere colored by cell type densities.

F) UMAP from E showing the number of microglia, without adjusting for total number of cells, in each plaque niche for the CosMx dataset.

G) Blue: correlation of estimated microglial densities (x-axis) in each Ssq plaque niche and the expression level of DAM marker genes in the plaque niche (DAM score, y-axis). Orange: correlation of estimated microglial densities (x-axis) and the expression level of HM marker genes (HM score, y-axis). Microglial densities were estimated using cell2location. DAM and HM scores were obtained by scoring the pseudo-bulked transcriptomes of the plaque niches using the *score_genes()* function in SCANPY. Ssq p-DAM scores for each plaque niche were calculated by subtracting the HM score from the DAM score.

*P < 0.1; **P < 0.01; ***P < 0.001

*Supplementary Figure 3:*

A) Violin plots comparing the raw microglial numbers in plaque niches across different brain regions analyzed in the CosMx dataset. One-way ANOVA followed by Tukey HSD multiple comparisons correction.

B) Violin plots comparing the raw astrocyte numbers in plaque niches across different brain regions analyzed in the CosMx dataset. One-way ANOVA followed by Tukey HSD multiple comparisons correction.

C) Pearson’s correlation analysis between the estimated microglial density (x-axis) in a plaque niche and its plaque DAM (Ssq p-DAM) level (y-axis) in Stereo-seq 3M (blue) and 18M (orange) plaque niches.

D) Violin plots comparing Ssq p-DAA levels across different brain regions in the two different ages (left: 3M, right: 18M). One-way ANOVA followed by Tukey HSD multiple comparisons correction.

E) UMAP of the Stereo-seq plaque transcripts data (from D) showing the expression level of certain cell type markers for each cell to decipher whether specific cell types drive the different clusters. Cell type markers were obtained from the Allen Brain Institute reference atlas (Yao 2021).

WM: white matter, ISO: isocortex, HIP: hippocampus, BS: brainstem; OLF: olfactory cortex; CTXsp: cortical subplate; *P < 0.1; **P < 0.01; ***P < 0.001; ns, not significant

*Supplementary Figure 4:*

A) MA plot showing significantly (p.adj < 0.05) up- (n = 29) and down-regulated (n = 326) genes in 3M Stereo-seq plaque niches with respect to p-DAM levels. Significantly changed genes are labeled in red. Quasi-likelihood F-test (QLFTest), P-values adjusted with Benjamini-Hochberg correction (P.adj < 0.1).

B) MA plot showing significantly (p.adj < 0.05) up- (n = 12) and down-regulated (n = 133) genes in 3M Stereo-seq plaque niches with respect to p-DAA levels.

C) Quadrant plot comparing gene expression changes in 18M plaque niches in response to Ssq p-DAM levels (y-axis) and to Ssq p-DAA levels (x-axis). 179 genes that were significantly altered in both conditions were located in Q1, 65 genes in Q2, 544 genes in Q3 and 24 genes in Q4. Red: genes significant in both comparisons, gray: genes not significant in either comparison, blue: genes significant in p-DAA comparison only, green: genes significant in p-DAM comparison only. Quasi-likelihood F-test (QLFTest), P-values adjusted with Benjamini-Hochberg correction (P.adj < 0.05).

D) MA plot showing absence of significantly (p.adj < 0.1) changed genes in CA1 neurons with respect to CosMx p-DAA levels.

E) MA plot showing significantly (p.adj < 0.1) down-regulated genes in CA1 neurons with respect to CosMx p-DAM levels.

F) MA plot showing significantly up-and down-regulated genes in DG neurons with respect to CosMx p-DAA levels.

G) MA plot showing significantly (p.adj < 0.1) up-and down-regulated genes in DG neurons with respect to CosMx p-DAM levels.

*Supplementary Table 1:*

Enrichment of GO Biological Processes terms in plaque niches with respect to Ssq p-DAM and Ssq p-DAA scores. Gene Set Enrichment Analysis (GSEA) results for both axes are listed in the xlsx file. NES = Normalized Enrichment Score

## Methods

### RESOURCE AVAILABILITY

#### Lead contact

Further information and requests for resources and reagents should be directed to and will be fulfilled by the lead contact, Professor Bart de Strooper.

#### Materials availability

This study did not generate new unique reagents.

#### Data and code availability

The spatial data will be deposited and will be publicly available as of the date of publication. Microscopy images will be shared by the lead contact upon request. Further information and requests for data analysis and data availability should be directed to and will be fulfilled by Professor Mark Fiers. The data are available from our website. Any additional information required to re-analyze the data reported in this paper is available upon request.

### EXPERIMENTAL MODEL AND STUDY PARTICIPANT DETAILS

All animal experiments were conducted in line with approved protocols by the Ethical Committee of Laboratory Animals of the KU Leuven. Male mice containing the *App^NL-G-F^* knock-in mutation^20^ expressed *App* mutations in the C57BL/6J background. *App^NL-G-F^* mice and wildtype (WT) controls were sacrificed at 3 months (3M) and 18 months (18M) using a carbon dioxide overdose. Following cervical dislocation, left and right hemispheres were embedded in cold OCT and snap-frozen in isopentane chilled with liquid nitrogen and stored at -80°C.

### METHOD DETAILS

#### Stereo-seq: experimental method

For Stereo-seq, 10 µm thick tissue sections were adhered to the Stereo-seq chip and incubated at 37°C for 3 minutes, with immediately adjacent slides collected for staining. Following the previously described protocol^18^, the sections were fixed in methanol for 30 minutes at -20°C. The sections were then stained for ssDNA and imaged to obtain nuclei location information. Tissue sections were then permeabilized for 12 minutes and reverse transcription performed overnight at 42°C. Afterwards, tissue was digested and cDNA released from the chip. The recovered cDNA was amplified and a total of 20ng was fragmented for library construction. PCR products were purified and sequenced on a MGI DNBSEQ-Rx sequencer. Two mixed hemispheres, where one hemisphere is from a WT mouse and the other from an *App^NL-G-F^* mouse, were processed for 3M animals, whilst two full coronal sections were processed for the 18M *App^NL-G-F^* mice, with one full coronal section of an age-matched control.

#### CosMx: experimental method

A previously described protocol was used for data acquisition^13^. Briefly, 10 µm thick tissue sections were fixed, antigen retrieval was performed and the tissue was permeabilized. In situ hybridisation was performed with the RNA mouse neuroscience panel, including the 18s RNA probe, and antibody morphology stain was performed with antibodies for Histone, glial fibrillary acidic protein (GFAP), 4′,6-diamidino-2-phenylindole (DAPI) and MOAB-2 for amyloid-β. Information from 27 FOV, with the size of 0.985 mm x 0.657 mm, were selected. After data acquisition, decoding of individual transcripts and segmentation of cells, based on Cellpose^43^, was performed as previously described^13^. Data was acquired from a coronal section of 18M mixed hemispheres of an *App^NL-G-F^* mouse and an age-matched control.

#### IHC staining of adjacent sections

Adjacent slides from the Stereo-seq experiments were stained using a standard IHC protocol to visualize reactive astrocytes and amyloid-β plaques. In brief, after fixation with 4% paraformaldehyde for 10 minutes and 90 minutes incubation with a blocking solution (5% donkey serum and 0.5% Triton X, in PBS) at room temperature, the tissue sections were incubated with GFAP antibody and anti-amyloid-β 3-8 (6E10) antibody conjugated with fluorophore Alexa 488 overnight at 4°C. The following day, sections were incubated with appropriate secondary antibodies (Goat anti-Guinea Pig Alexa 568) for 90 minutes at room temperature. After five wash steps, autofluorescence was quenched by treating sections for 30 seconds. with 1x TrueBlack solution diluted in 70% ethanol. Afterwards, the sections were rinsed by dipping the slides 20x in PBS. Nuclei were visualized by incubating the sections with DAPI staining for 15 minutes at room temperature. Sections were rinsed in SSC buffer, before 100 µl glycerol was added and a coverslip was applied.

16-bit fluorescent images of DAPI, 6E10, and GFAP were acquired on a Zeiss Axioscan.Z1 slidescanner, with a 20x/0.8 NA air objective using the ZEN blue software (version 3.1, Carl Zeiss Microscopy GmbH). Different FOV were stitched together using the online stitching configuration in the software.

### QUANTIFICATION AND STATISTICAL ANALYSIS

#### Stereo-seq: data processing

The Stereo-seq raw data processing was performed as described^18^ and in the publicly available SAW pipeline (https://github.com/BGIResearch/SAW). In brief, fastq files were generated using a MGI DNBSEQ-Tx sequencer. CID sequences (1-25 bp) on the first reads were first mapped to the designed coordinates of the in situ captured chip, allowing 1 base mismatch. Low quality reads were filtered out, and retained reads were aligned to the mm10 reference genome using STAR^44^. Mapped reads with MAPQ > 10 were counted and annotated to their corresponding genes. UMIs with the same CID and the same gene locus were collapsed, again allowing 1 base mismatch, resulting in a CID-containing expression profile matrix.

#### Image-based single cell segmentation

Single cell segmentation was performed by first aligning the nucleic acid staining to the Stereo-seq chips and then applying the watershed algorithm through the Scikit-image package (V0.18.1). The number of markers required for the watershed algorithm were obtained through Gaussian-weighted local threshold binarization with block size of 41 and offset of 0.003. We then exacted Euclidean distance transformation (with distance of 13 or 15) from the background removed images. For each of the segmented cells, UMIs from all DNB within the corresponding segmentation were aggregated per-gene and then summed to generate a cell by gene matrix. The centroid of each cell was determined using rearrr (https://github.com/LudvigOlsen/rearrr).

#### Stereo-seq raw data processing

Gem files specifying the X/Y coordinates and MID count for every DNB were read in using stereopy’s read_gem() (https://github.com/BGIResearch/stereopy) to generate both a bin20 expression matrix (with argument bin_size = 20) and a cell_bin expression matrix (argument bin_type = ‘cell_bins’). Both datasets were manually aligned with 13 brain regions defined by the Allen Brain Mouse Atlas^21^, and each cell/bin was assigned to a region (Supplementary Fig 1A). Cells/bins overlapping with white matter and with the cerebral nuclei region were removed. The 13 brain regions were further grouped into 5 coarse regions (brain stem, cortical subplate, hippocampus, isocortex, and olfactory bulb). To then remove low-quality bins, bins with a high or a low number of genes, MIDs, mitochondrial content were removed. Furthermore, genes expressed in less than 0.1% of bins in a given sample were removed.

#### Transcript visualization

Visualization of spatial localization of Stereo-seq transcripts, overlaid over segmented plaque images, was performed using TissUUmaps^45^.

#### CosMx: data processing

The expression matrix, metadata, and FOV files were read in using squidpy’s *read_nanostring()* function. Transcripts associated with negative and custom probes were immediately removed from the analysis. Cells were manually assigned regional annotation based on visual alignment with the Allen Brain Mouse Atlas reference (Supplementary Fig 1G). Cells with a high or low number of genes or transcripts were removed. In order to identify likely mis-segmented doublets (i.e. two cells that were incorrectly identified as one cell during the segmentation process), scrublet^46^ was run on the AD and the WT hippocampi, separately. Cells that were assigned a high doublet probability were removed. Initial clustering of the dataset revealed a unique cluster of cells that all overlapped with plaques and expressed aberrantly high levels of a set of transcripts not known to be associated with amyloid pathology (*Rims1, Drd4, Lilra5,* among others). Upon further analysis, it was concluded that the probes associated with these transcripts likely bind nonspecifically to amyloid plaques. The relevant cluster of cells was therefore removed from subsequent analysis.

#### Unsupervised clustering of the cells and bins

Cells/bins passing quality control in each of the datasets were further processed using the SCANPY workflow. Raw gene expression counts were first log-transformed and normalized by library size using the *normalize_total()* and *log1p()* functions, and subsequently scaled to unit variance and zero mean with the *scale()* function. Principal Component Analysis (PCA) was performed on the expression profile of the genes. Cells/bins from different Stereo-seq chips were integrated using Harmony^47^ to account for inter-chip batch effects. The first 20 PCs were used as input for dimensionality reduction with Uniform Manifold Approximation and Projection (UMAP). CosMx cells were clustered using the Leiden method at a resolution of 0.8 to obtain 11 clusters. Clusters were annotated for cell type based on relative expression of cell type marker genes, (determined with the *score_genes()* function). The subclusters characterizing the glutamatergic neuron cluster were further annotated based on their regional spatial distribution.

#### Amyloid segmentation

Amyloid-β plaques were annotated using the QuPath software through a pixel classification threshold with an additional size threshold of 65,000 px to discriminate the pathology from background and artificial staining. The segmentation was then manually checked to ensure segmented plaques were of high quality. The segmented plaques were then exported to ImageJ and converted into a binary mask for alignment to the transcriptome.

Using a previously described technique^8^, the adjacent slide staining was aligned to the transcriptome of the Stereo-seq chip. In short, corresponding landmarks were selected on both the staining and the transcriptome. Using the Fiji “Landmark correspondences” plugin, the staining was then aligned to overlay with the transcriptome. The same method was followed for alignment of the Allen Brain Atlas with the transcriptome that allowed for assignment of brain regions to the transcriptome. After alignment of adjacent slide stainings, segmented masks from either side of each chip were summed to obtain one binary amyloid segmentation mask per chip.

#### Calculation of distances to plaques

The distance from the center of each bin/cell to the edge of the nearest segmented plaque was calculated using Scipy’s KDTree package: (https://github.com/scipy/scipy/blob/main/scipy/spatial/kdtree.py).

#### Dimensionality reduction and deconvolution of Stereo-seq plaque niches

The transcriptomic profiles of plaque niches in the Stereo-seq dataset were acquired by “pseudo-bulking” all the transcripts identified within 40 µm (80 pixels) from the edge of a plaque, inclusive of those overlaying the plaque. Plaque niches with less than 10 transcripts, as well as lowly expressed genes, were discarded from further analysis. The resultant expression matrix was log-normalized and total transcript counts were regressed. Principal Component Analysis (PCA) was performed, and the niches were integrated across samples using Harmony^47^. The first 10 principal components were used to construct the UMAP and a resolution of 0.3 was used to cluster the niches. Cell2location was then used to infer the cellular composition of these pseudo-bulked plaque niches, using the scRNA-seq dataset from the Allen Brain Mouse Atlas^23^ as the reference dataset.

#### Plaque niche neighborhood analyses

Plaque x cell type density matrices were created by counting, for every plaque, the number of each given cell type lying within a 40 µm distance (80 pixels in Stereo-seq, 240 pixels in CosMx) from the plaque’s edge and dividing that number by the total number of cells within that radius. In the case of the CosMx dataset, segmented cells were counted, only taking plaques into consideration that contained 5 or more cells. For Stereo-seq, cell type compositions inferred by cell2location were used. The resultant matrices were used for dimensionality reduction.

#### p-DAM and p-DAA score calculation

To obtain p-DAM and p-DAA scores of plaque niches in Stereo-seq (Ssq p-DAM and Ssq p-DAA), pseudo-bulked plaque niche transcriptomes (*i.e.* all transcripts identified within 40 µm of a plaque edge, as described in “*Dimensionality reduction and deconvolution of Stereo-seq plaque niches”)* were normalized and scored on their relative expression of DAM marker genes, HM marker genes^3^, DAA markers, and homeostatic astrocyte markers^5^ using SCANPY’s *score_genes()* function. The Ssq p-DAM score was calculated as DAM score minus HM score, while the Ssq p-DAA score was calculated as DAA score minus homeostatic astrocyte score. The CosMx p-DAM and CosMx p-DAA scores were obtained by taking the max DAM score of any microglia and the max DAA score of any astrocyte, respectively, in a given plaque niche.

#### Differential gene expression analyses

DE analyses were conducted by fitting generalized linear models (GLM). Each GLM model was tested for DE with EdgeR’s quasi-likelihood F-test (QLFTest) which accounts for the uncertainty in dispersion estimation. The differential expressions were performed with respect to the plaque niche activation measure as a continuous variable. DEs in the CosMx dataset were performed on individual cells and a significance threshold of p.adj < 0.1 was used. When performing DEs on plaque niches in the Stereo-seq dataset, we controlled for the brain region and the estimated abundance of each cell type in the plaque niche (as estimated by cell2location). In these DEs, a significance threshold of p.adj < 0.05 was used.

#### Gene Set Enrichment analyses

Gene set enrichment analyses (GSEA) were performed using R’s clusterProfiler package^48^, using the log2 fold change value for gene rank. In the quadrant analysis of enriched GO terms, GO terms in the same quadrant were grouped together semantically using the rrvgo package^49^. The average NES and minimum FDR values of the subterms were taken as the parent term’s NES and p-value.

#### Cell type reference

The 2020 10X Genomics scRNA-seq dataset for the mouse whole cortex and hippocampus was downloaded from the Allen Brain Atlas Institute webpage. For analysis, a subset of the data was generated to only include regions annotated for ’RSP’, ’TEa-PERI-ECT’, ’ACA’, ’AI’, ’SSs-GU-VISC-AIp’, ’AUD’, ’MOp’, ’MOs_FRP’, ’PL-ILA-ORB’, ’PTLp’, ’SSp’, ’VIS’, ’VISl’, ’VISm’, ’VISp’, ’HIP’. The remaining cells were randomly sampled to retain 12.5% of the cells (131,169 total cells). Wilcoxon rank sum test was used to obtain the top 200 markers for annotated cell types (Glutamatergic neurons, GABAergic neurons, microglia, astrocytes, endothelial, and oligodendrocytes).

