## Supplementary figures and images for "Applying high-resolution spatial transcriptomics to characterize the amyloid plaque cell niche in Alzheimer’s Disease"

### Supplemental Figure 1

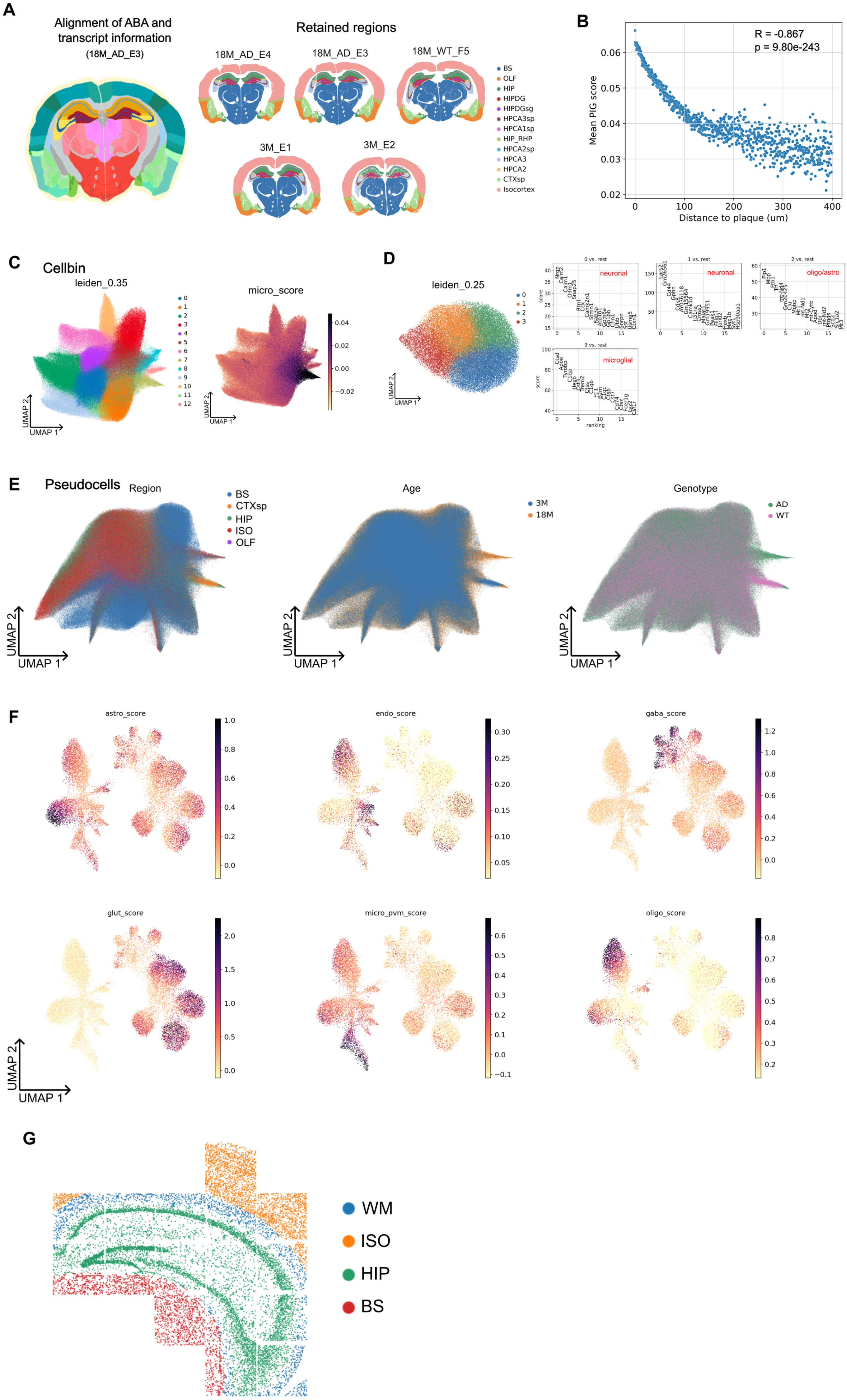

### Supplemental Figure 2

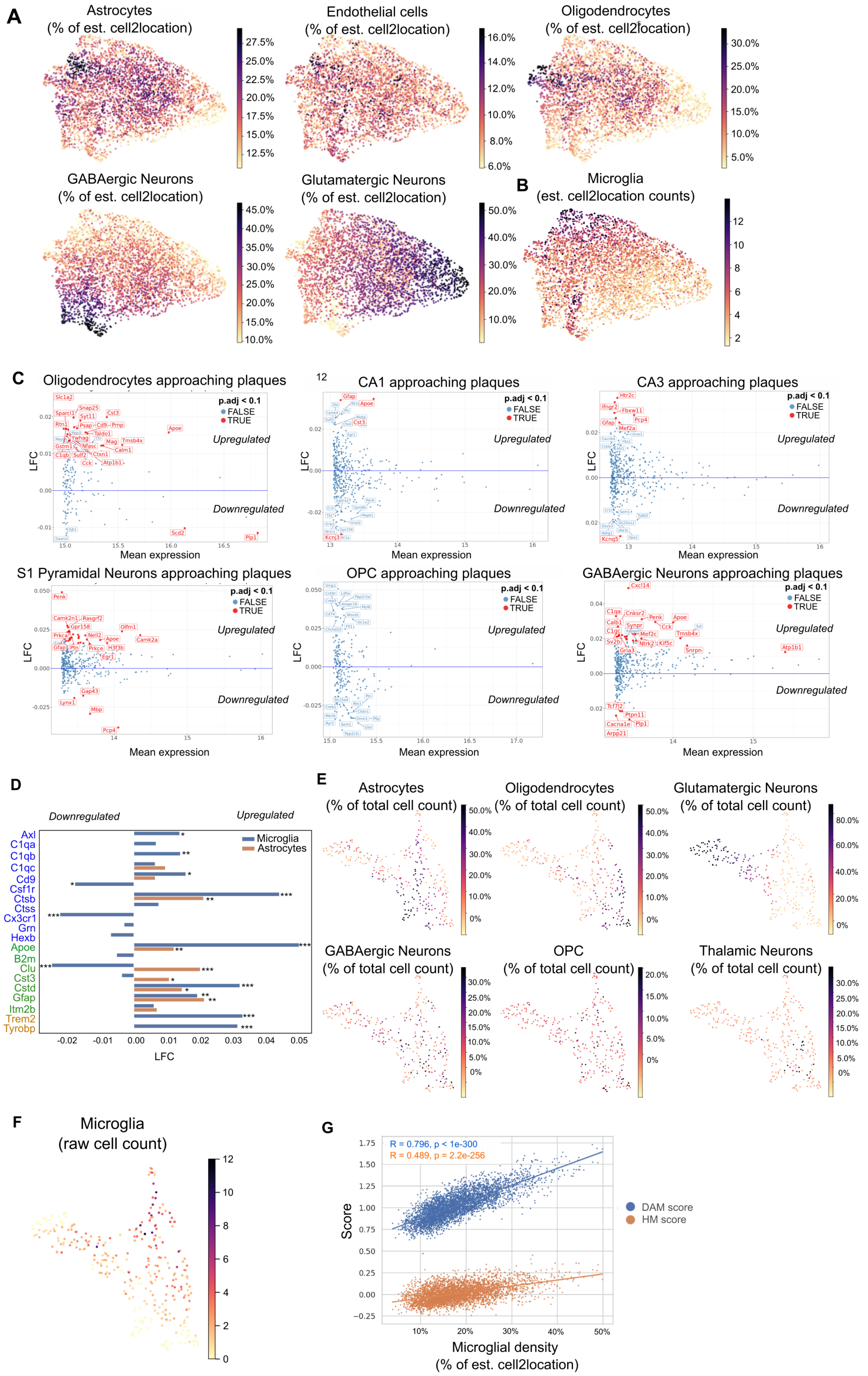

### Supplemental Figure 3

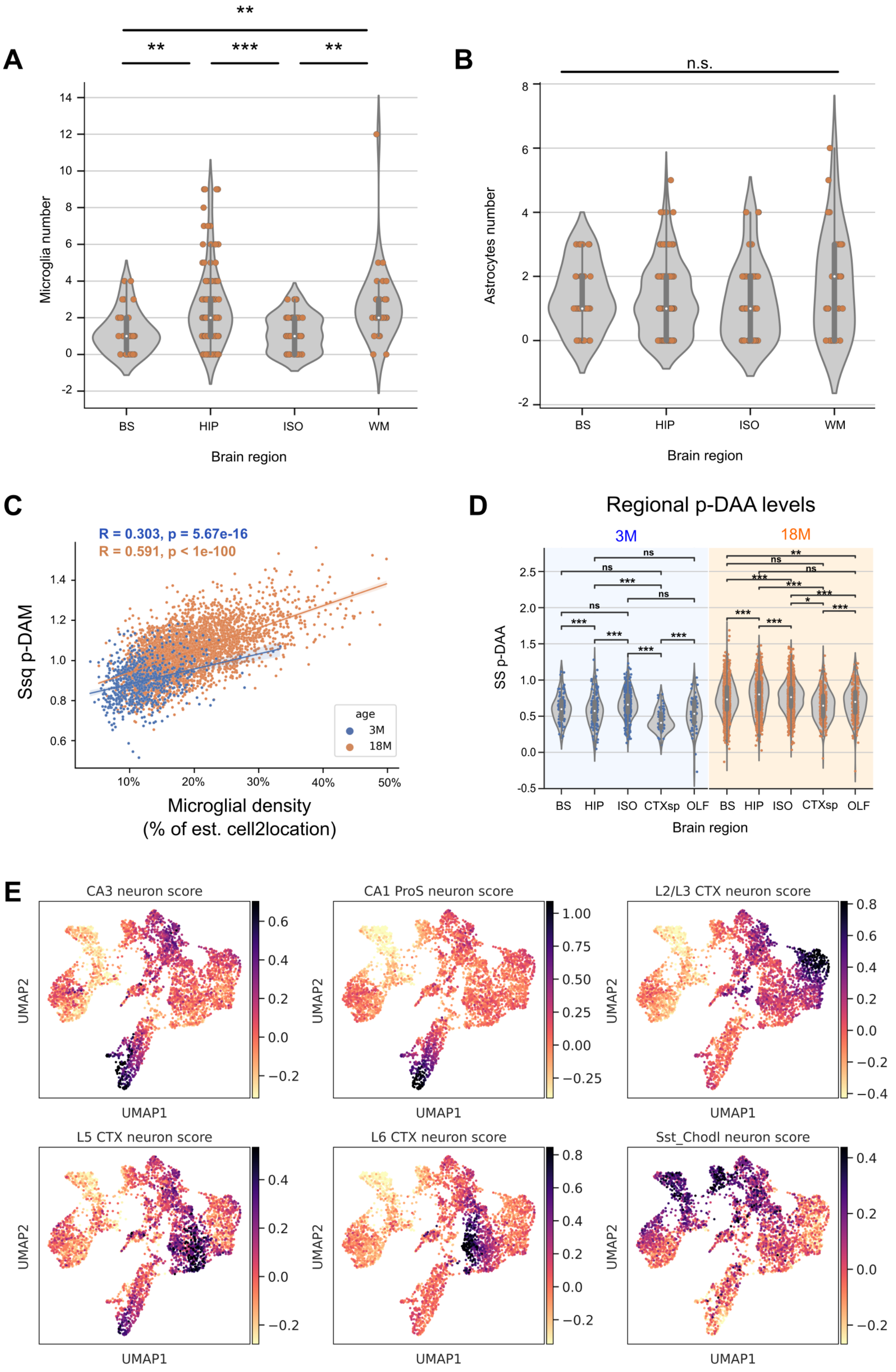

### Supplemental Figure 4

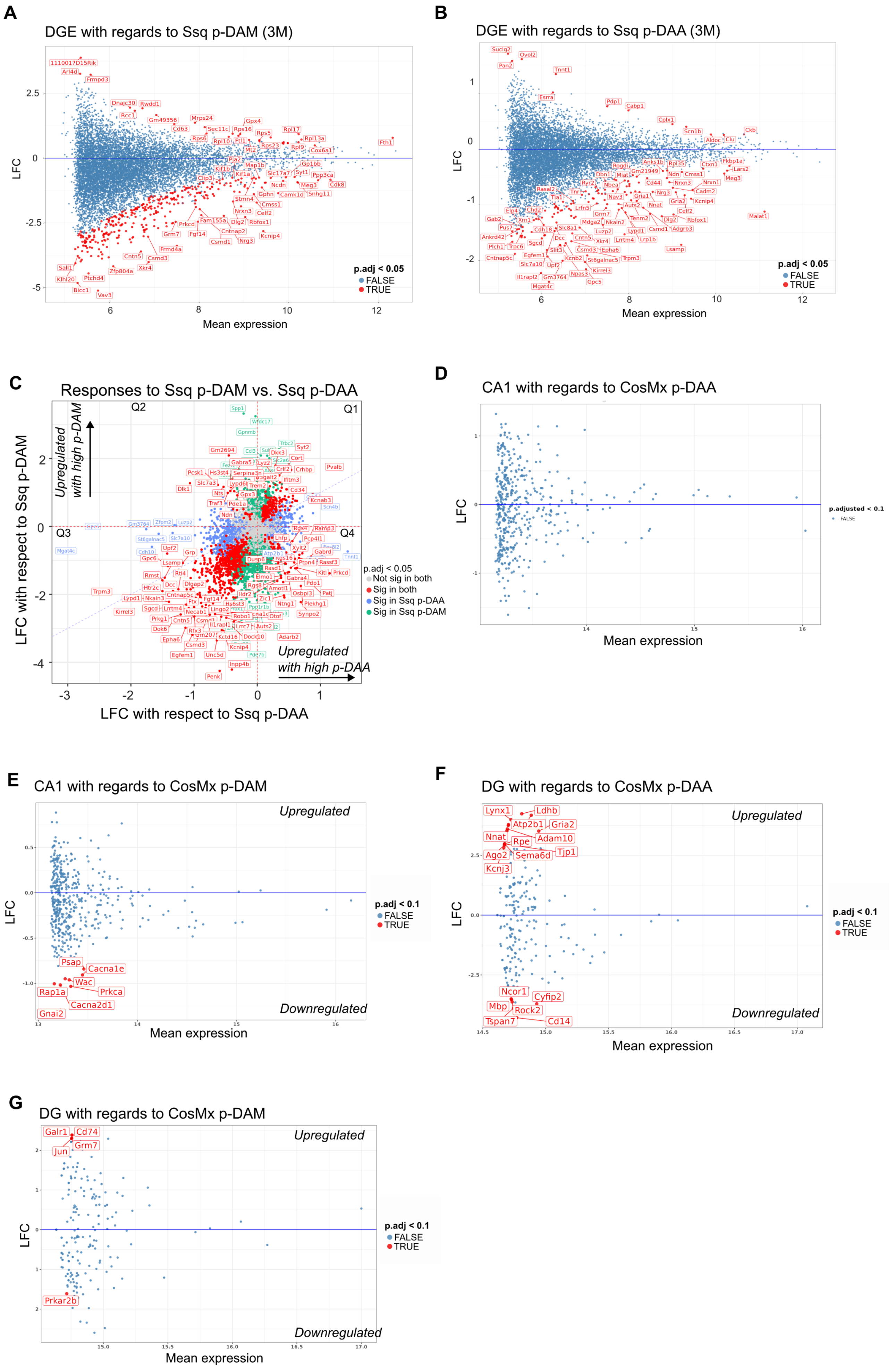
